# Whole-bone toughness is linked to canal and osteocyte lacunae deficits in the ZDSD type 2 diabetic rat model

**DOI:** 10.1101/2023.03.07.531548

**Authors:** William Woolley, Yoshihiro Obata, Kaitlynn Martin, Claire Acevedo

## Abstract

Type 2 diabetes mellitus (T2DM) is associated with an increased fracture risk independent of bone mass. The exact origin of this increased fracture risk is still not well understood. Using a polygenic diabetic rat model, synchrotron radiation micro-computed tomography (SRμCT), and *in situ* scanning electron microscope (SEM) fracture toughness, we related the changes at the microscale to toughness and material properties of diabetic rat femurs. The diabetic rat model (ZDSD) displayed overnight fasting hyperglycemia and an increased AGEs content. Additionally, we measured the impairment of post-yield properties and toughness in diabetic rats. The cortical geometry and porosity were also affected in this ZDSD model. We measured a decrease in osteocyte lacunar density associated with a decreased lacunar volume. Moreover, we found decreased canal density while maintaining a similar canal diameter. These results indicate that diabetes impairs bone remodeling, affecting bone microstructure. Because canals and lacunae are also linked with extrinsic toughening mechanisms, we attribute the decreased toughness largely to these microstructural changes. In conclusion, we showed that changes in lacunae and canal density, combined with AGEs accumulation, decreased toughness in T2DM rat bone.

## Introduction

In 2015, the prevalence of diabetes among adults was estimated to be 8.8%, and based on the Centers for Disease Control and Prevention, approximately 90-95% of diabetes cases are type 2 diabetes mellitus (T2DM). In 2040, these numbers are projected to increase to 10.4%, reaching global epidemic levels [1]. T2DM is correlated with an increased bone fracture risk for a given bone mass [2–4], which can lower life expectancy [1, 5]. The decrease in fracture resistance induced by diabetes has been well established [6–10], but the exact origin of this decrease is still not well understood. Gaining insights into the origins of bone fragility in diabetic patients would improve the prediction, prevention, and treatment of such a fast-growing disease.

In T2DM, hyperglycemia and oxidative stress boost the formation of non-enzymatic cross-links, also called advanced glycation end-products (AGEs), via the Maillard reaction [11]. Normally, AGEs accumulate with age at a slow rate, but with T2DM, the accumulation rate is increased [12]. AGEs accumulation is a well-known contributor to increased fracture risk [13, 14]. Mechanisms by which AGEs alter the whole-bone behavior in T2DM have been partially revealed through collagen stiffening and impairment of collagen’s ability to deform, restricting post-yield bone properties [15–19]. In addition, AGEs accumulation is also thought to impair osteoblast, osteoclast, and osteocyte cellular function [20–24], disrupting bone remodeling, which is essential to maintain bone quality and resistance to fracture. In particular, osteocytes - the most abundant bone cells regulating bone remodeling and bone homeostasis-are altered when exposed to AGEs and hyperglycemia *in vitro* [25]. Disruption of osteocyte-mediated remodeling might cause distinctive changes in bone microstructure, osteocyte lacunar density, osteocyte lacunar size, and osteocyte apoptosis [26, 27]. In T2DM, microstructural changes were reflected in increased cortical porosity (i.e., the ratio of pore volume to bone tissue) [10]. There is a need to investigate whether increased cortical porosity can be explained at the microscale by changes in osteocyte lacunae and canals using a diabetic animal model that closely mimics diabetes in humans.

The exact effect of T2DM on bone geometry and microstructure is not well understood. For example, some studies report a decreased cortical thickness and a reduced cortical area [28, 29], whereas in others, the cortical area is similar between control and diabetic patients [30]. At the microscale, studies consistently report that cortical porosity is increased with diabetes [9, 10], and cortical porosity has been shown to influence the crack path and fracture resistance [31–33]. Because canals and lacunae are inherently linked to porosity and fracture in bone, these features are also of chief interest. Changes in bone geometry will influence the force and displacement that the bone can sustain but will not affect the bone properties expressed in terms of stress and strain. Changes in bone microstructure have the potential to alter the crack path and the crack-growth toughening mechanisms [34].

Fracture resistance in bone originates from two main types of toughening mechanisms: intrinsic and extrinsic [35, 36]. Intrinsic (material) toughening mechanisms occur at the nanoscale-collagen level, with salient phenomena such as fibrillar sliding granting bone its enhanced plasticity and toughness [37, 38]. Intrinsic toughening mechanisms act at the tip of the crack to prevent crack initiation and growth through collagen-induced plasticity and can thus be restricted by AGEs accumulation [15, 39, 40]. Extrinsic toughening mechanisms are largely conferred by microstructural features; these mechanisms influence crack-growth toughness [41]. In bone, extrinsic toughening mechanisms act to arrest crack growth by acting in its wake [35]. Features of a propagating crack such as a tortuous crack path or crack bridging work to dissipate energy during crack growth [41]. For extrinsic toughening mechanisms, osteons and their cement lines are key features of bone microstructure for deflecting a crack and for dissipating the energy of crack propagation [42]. Since canals lie in the center of the osteons, their number and size may provide insight into changes in crack-growth toughness.

To understand how T2DM alters microstructural features, we used high-resolution synchrotron radiation micro-computed tomography (SRμCT) to reveal the internal composition of bone at the microscale, including osteocyte lacunae, canals, and mineral content. In this paper, we tested the hypothesis that T2DM significantly impacts osteocyte lacunae and osteonal canals, which will impair bone quality via remodeling and crack-growth resistance. To test this hypothesis, we chose a Zucker Diabetic Sprague Dawley (ZDSD) rat model and compared toughness, material properties, geometry, and microstructure to age-matched control rats. Specifically, we measured toughness by performing *in situ* scanning electron microscope (SEM) fracture toughness tests, tissue material properties by performing strength tests, and geometry and microstructure, using optical and SRμCT imaging. For the first time, we quantitatively relate lacunae and canals to the toughness and material properties of diabetic bones.

## Materials and Methods

### Study design

This study was designed to evaluate the effect of T2DM on geometry and microstructural features and relate them to whole-bone toughness. We used a well-established ZDSD rat model that mimics the pathophysiology of human T2DM with polygenic phenotype, intact leptin pathway, glucose intolerance, and hyperglycemia [10, 43]. This rat model develops diabetes at an adult age (≥16 weeks) with a pre-diabetic phase (8-16 weeks). All rats used in this work were male rats. We used an age-matched, non-diabetic control rat model, lean Sprague Dawley (LSD), of the same sex to compare against the ZDSD rats. In the current work, we aimed to measure how the ZDSD rat model influenced (1) bone tissue mechanical properties, including toughness, and (2) features of bone microstructure, including canals and osteocyte lacunae.

### Animals and tissues

Nineteen-week-old ZDSD male rats (‘‘diabetic”; n=20, mean weight = 478 g) and LSD (‘‘control”; n=20, mean weight = 518 g) were purchased from Charles River and used in this study. Rats were maintained and treated in accordance with Institutional Animal Care and Use Committee (IACUC)-approved protocols. The generation and phenotypes of these rats have been described previously [44]. Rats were fed with standard chow (Catalog No. 2920x; Envigo Teklad Global). We withdrew blood by intra-cardiac punctures at the time of sacrifice to measure circulating glucose after an overnight fast. Left femora were harvested from the rats. After removing soft tissues, the proximal and distal epiphysis were cut with a low-speed diamond-blade saw under irrigation to isolate the diaphysis. Femora were stored in phosphate-buffered saline (PBS) at 4°C for mechanical testing.

### Biochemical quantification of AGEs

Using a low-speed diamond-blade saw, approximately 30 mg of cortical bone was cut from the femur used in the mechanical test and demineralized in 10% ethylenediaminetetraacetic acid (EDTA) for seven days. The organic matrix was digested by proteinase K (1 mg/ml) in a 60°C water bath for three hours. Fluorescence was read using a microplate reader (SpectraMax M2, Molecular Device, USA) at 370/440 nm excitation/emission and compared with a quinine standard to obtain the AGEs content. AGEs content was then normalized to the collagen content by performing the hydroxyproline assay as described by Brown et al. [45]. Briefly, hydroxyproline is oxidized to a pyrrole which then reacts with p-dimethylaminoazobenzene (p-DAB), allowing the solution to be read by absorbance at 561 nm.

### Flexural strength test

To assess flexural mechanical properties, we tested femora (n=5-8/group) in three-point bending with the load placed on the anterior side. This setup placed the anterior side of the rat femurs in compression and the posterior side in tension. The test was performed in a Psylotech MicroTest System (Psylotech Incorporated, Evanston, IL, USA) under displacement control, with a 220 N load cell. During these three-point bending tests, the span was 25.4 mm, and the displacement speed was 833 nm/s. All bone samples were wrapped in PBS-soaked gauze to maintain sample hydration during testing. The procedure of the strength test was performed in accordance with ASTM D790 [46]. Force and displacement data were recorded during testing and were then used to calculate stress and strain using beam theory [47]. Bone properties (bending modulus, ultimate stress, ultimate strain, strain to failure, work to fracture) were then evaluated using geometric parameters obtained from optical microscopy. All calculations were performed in Python.

### *In situ* SEM fracture toughness test

To evaluate fracture toughness, we performed three-point bending on single-edge notched femora (n=6-7/group) through an *in situ* SEM fracture toughness test to measure resistance to fracture while simultaneously imaging crack extension. Samples were notched on the posterior side using a custom-built razor notcher and a 1 μm diamond suspension, providing an initial crack size of approximately one-third of the anterior-posterior diameter with a tip radius of ≤ 5 μm. The notch dimensions were made in accordance with ASTM E1820 recommendations and the method explained by Ritchie et al.[47, 48]. Fracture toughness tests were performed in three-point bending (8 mm loading span) on PBS-soaked samples at 25°C under displacement control, at 833 nm/s, using a Psylotech MicroTest System (Psylotech Incorporated, Evanston, IL, USA) placed inside a low-vacuum SEM (JEOL JSM-5910LV). The pressure was set to 50 Pa to maximize the image quality while maintaining hydration in the bone. During the test, the loading was paused after each crack increment to record the load and crack length. Images of the crack path were captured with the back-scattered electron mode at a voltage of 25 kV.

We used crack-resistance curves, or R-curves, to capture the contribution of plastic deformation and crack growth in fracture toughness, represented by the K-equivalent stress intensity factor, *K_eq_* [47]. First, the nonlinear strain-energy release rate, *J*, was obtained from the sum of the elastic contribution, *J_el_*, based on linear-elastic fracture mechanics in mode I, and the plastic contribution, *J_pl_*, for a stationary crack in bending. *K_eq_* values were back-calculated from the *J* measurements using the standard J-K equivalence for nominal mode I fracture, specifically that *K_j_* = (*JE*)^1/2^, using the bending modulus values measured during strength tests. To ensure K-dominance at the crack tip, crack length (*a*) should be roughly an order of magnitude smaller than the bone diameter, in our case: *a* <450-500 μm [47]. All calculations were performed in Python.

### Light microscopy for bone geometry measurement

Following strength and toughness tests, we captured the fracture surface using an Olympus SZX16 microscope equipped with a 14 MP microscope camera (AmScope MU1403). We imported the images into ImageJ (Fiji) [49] to measure geometrical parameters. First, we created a binary mask of the bone by applying a pixel intensity threshold. Then, using BoneJ (version 1.4.3.) [50], we measured the cortical and medullary areas, the cortical thickness, and the moment of inertia in the anterior/posterior and medial/lateral axes. Using the binary bone mask, we fit ellipses to the endosteal and periosteal surfaces of the bone to measure their radii. Finally, we measured the notch angle representing the initial angle of the notch during toughness testing.

### Synchrotron radiation micro-computed tomography

Microscale imaging was performed at the synchrotron microtomography beamline 8.3.2 at the Advanced Light Source in Berkeley, CA. Images were acquired with a beam energy of 18 keV and a 250 ms exposure time. A total of 1969 projection angles were acquired over a 180° sample rotation. After image acquisition, reconstruction of SRμCT images was performed in open-source Python code TomoPy [51]. The final image pixel spacing was 1.6 μm/pixel. Four control and six diabetic bone samples from the *in situ* SEM fracture toughness test were imaged.

### Synchrotron radiation micro-computed tomography image analysis

Mineralization, lacunae properties, and canal properties were analyzed to compare the differences between the control and diabetic bones in this study. All analyses used a bone volume with a depth of 300 image slices (480 μm). For multiple analyses, a binary mask of cortical bone was necessary. This bone mask was created in Dragonfly 2020.2 (Object Research Systems (ORS) Inc., Montreal, Canada) by applying a pixel-intensity threshold to an 8-bit image of the bone, followed by closing holes smaller than 60 μm. Next, the image was inverted and a particle filter was applied to exclude all features other than the inverted bone, and then inverted back. This bone mask was used to calculate the tissue volume in the bone for the normalization of canal density and lacunar density and was used as an image operator for canal and mineralization calculation.

Assessment of bone mineral content was performed on the original 32-bit images in ImageJ (Fiji). First, the mineralization of the bone tissue was calculated by performing a multiplicative operation of the binary bone mask and the 32-bit bone image. Image gray values were then recorded using the image histogram calculator in Fiji. These gray values were converted to mgHA/cm^3^ by using the attenuation coefficient of hydroxyapatite in bone; this measure of hydroxyapatite is known as volumetric tissue mineral density (vTMD). Numerical analysis of vTMD peak and full-width half-max (FWHM) was performed in Python.

Quantifying lacunae was performed by first performing a median filter of radius 1 on the original 32-bit images using Fiji and adjusting the brightness and contrast uniformly. These images were then converted to 8-bit and imported into Dragonfly, where a common pixel intensity threshold was used to separate the background pixels from the bone. A particle size filter of between 8-600 voxels, approximately 18-1318 μm^3^, was then used to eliminate features larger or smaller than typical lacunae in rat bone. Data for all lacunae were exported from Dragonfly and imported into Python, where they were then binned by volume into a histogram for analysis of mean lacunar volume and lacunar density.

The diameter and density of canals were calculated by using the 8-bit bone images from the analysis of lacunae. A Gaussian blur filter with a radius of 2 pixels was applied to the images to obscure small features such as lacunae and noise. A pixel-intensity threshold was then used to segment the canals and background from the bone. The intersection of the bone mask and canal segmentation is then taken to isolate the canals. The BoneJ thickness tool was used in Fiji to calculate the local thickness of the canals. The number of canals was counted for each image slice and divided by the bone area in each image slice to calculate the volumetric canal density. All numerical analysis of canals was performed in Python.

### Statistical analysis

We performed independent t-tests using the SciPy package in Python [52] to determine group differences in terms of mechanical properties, geometry/microstructural features, crack-growth toughness, and AGEs content. For the mechanical tests, our sample size was 6-7/group for the toughness test and 5-8/group for the strength test. For the AGEs, our sample size was 5/group; for the SRμCT, the sample size was 4-6/group; for the geometry, the sample size was 12-15/group. Despite the small sample size per group, our data did not violate normality or equal variance assumptions (Shapiro-Wilkes test). Significance is defined by p<0.05, and data are given as mean ± standard deviation.

## Results

### The ZDSD rat model displays hyperglycemia and AGEs accumulation characteristic of T2DM

Blood glucose levels after overnight fasting were 44% higher in diabetic rats compared to aged-matched control rats (169.25 mg/dL vs 242.95 mg/dL, p<0.001, n=20/group) (Fig. 1a), which confirmed hyperglycemia in ZDSD rats. ZDSD rats display a progressive development of the disease with a pre-diabetic state (8-16 weeks), diabetic state (>16 weeks), and diabetic complications (>24 weeks). High blood glucose is indicative that despite being in an early diabetic phase, these rats have an altered insulin pathway. As expected from previous studies [15], high levels of circulating blood glucose significantly increased AGEs content in femurs. In diabetic rats, AGEs concentration per collagen content were 54% higher than in control rats (p<0.005, n=5/group) (Fig. 1b) [10]. In addition to displaying an altered insulin pathway in the early diabetic phase, ZDSD rats possess increased AGEs content at nineteen weeks of age compared to the control rats.

**Fig. 1.**
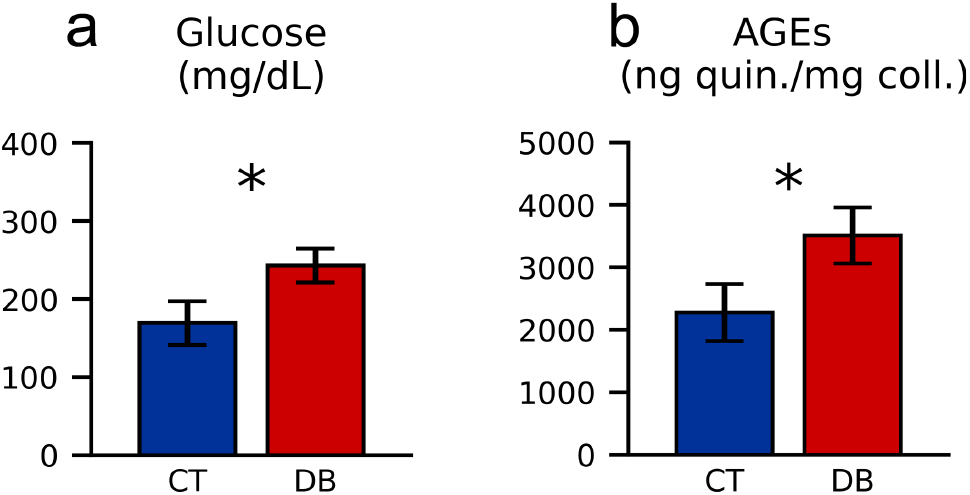
Blood glucose and AGEs quantification. (a) The overnight fasting blood glucose level was quantified in control (CT) and diabetic (DB) rats. Hyperglycemia was confirmed in diabetic rats with a 44% increase compared to controls (p<0.01, n=20/group). (b) The accumulation of AGEs was quantified using a fluorometric assay. These results indicate that the diabetic bones contained 54% more fluorescent cross-links than the control bones (p<0.01, n=5/group). Data are given as mean ± s.d. Groups were compared with an independent t-test.

### T2DM impairs post-yield properties and toughness

We performed flexural strength and *in situ* SEM fracture toughness tests to investigate the effects of T2DM on bone mechanical properties. We found significant changes in post-yield strength properties, with a 13% decrease in the strain to failure of ZDSD rat femurs, compared to control rat femurs (p<0.05, n=4-8) (Fig. 2f). Changes in strain to failure were reflected in work to fracture, decreasing the energy to failure by 14% in ZDSD rat femurs (Fig. 2g). Ultimate strength was not affected by the disease, with changes of −0.6% (p=0.81) (Fig. 2d). Additionally, no statistically significant changes were found in the pre-yield region, exhibited by the bending modulus (p=0.61) (Fig. 2c) and the yield strain and strength, with changes of −12% (p=0.25) and −12% (p=0.22), respectively. Fracture-toughness properties, presented as an equivalent stress-intensity, *K_eq_*, K-based crack-resistance curve (Fig. 2b), revealed a decrease in crack-growth toughness (slope of the R-curve) with diabetes. The diabetic rat femurs exhibited a 55% decrease in crack-growth toughness compared to the control rat femurs (p=0.09) (Fig. 2h).

**Fig. 2.**
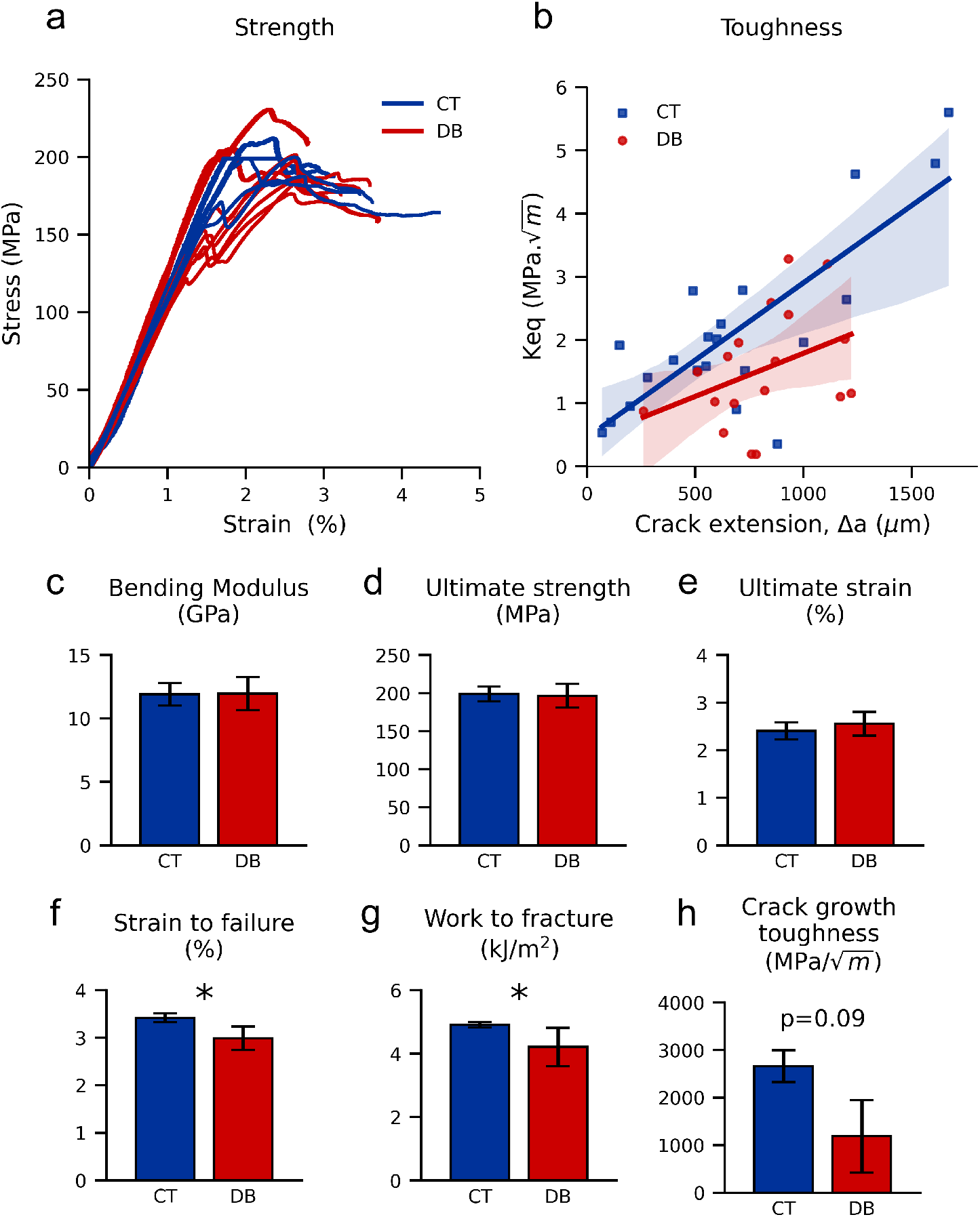
Flexural strength and *in situ* SEM fracture toughness test. Mechanical properties in rat femurs comparing control (CT) and diabetic (DB) groups are shown by (a) flexural strength (n=5 CT, n=8 DB), and (b) *in situ* SEM fracture toughness R-curve (n=6 CT, n=6 DB) properties. For flexural strength tests, (c) bending modulus, (d) ultimate strength, and (e) ultimate strain were not statistically significantly altered by diabetes, whereas (f) strain to failure (−13%, p=0.01) and (g) work to fracture (−14%, p=0.05) were significantly lower in the diabetic group than in the control group. For fracture toughness, (h) crack-growth toughness (slope of the R-curve) shows a lowered trend with diabetes (−55%, p=0.09). Data are given as mean ± s.d. Groups were compared with an independent t-test.

### T2DM reduces the whole-bone cortical geometry and porosity

Using microscopy (n=15 control and n=12 diabetic) and SRμCT (n=4 control and n=6 diabetic), we measured bone geometry, porosity, and mineralization parameters to assess how T2DM impacted bone shape and composition. In diabetic rats, the cortical bone area (Ct.Ar), the medullary cavity area (Me.Ar), and the cortical thickness (Ct.Th) are all decreased significantly by approximately 10% (Fig. 3b, c, f) in correlation with an 8% lower body weight in the diabetic animals. These reductions in bone geometry parameters affect the moment of inertia around both axes (Fig. 3d, e), reducing the resistance to bending, especially the resistance to moment about the medial-lateral axis. In addition to whole-bone geometry parameters, cortical porosity and vTMD were assessed using SRμCT. Using this technique, we found that cortical porosity is decreased (30% decrease, p=0.06) while the peak vTMD is maintained in diabetic rats’ femurs (Fig. 3g, h). The distribution of vTMD was altered, however. The FWHM of the vTMD distribution was significantly lower in the diabetic rat samples than in their age-matched controls (10% decrease, p=0.02) despite the unchanged vTMD peak, implying diabetic rats had a smaller variation of mineralization than their control counterparts (Fig. 3i).

**Fig. 3.**
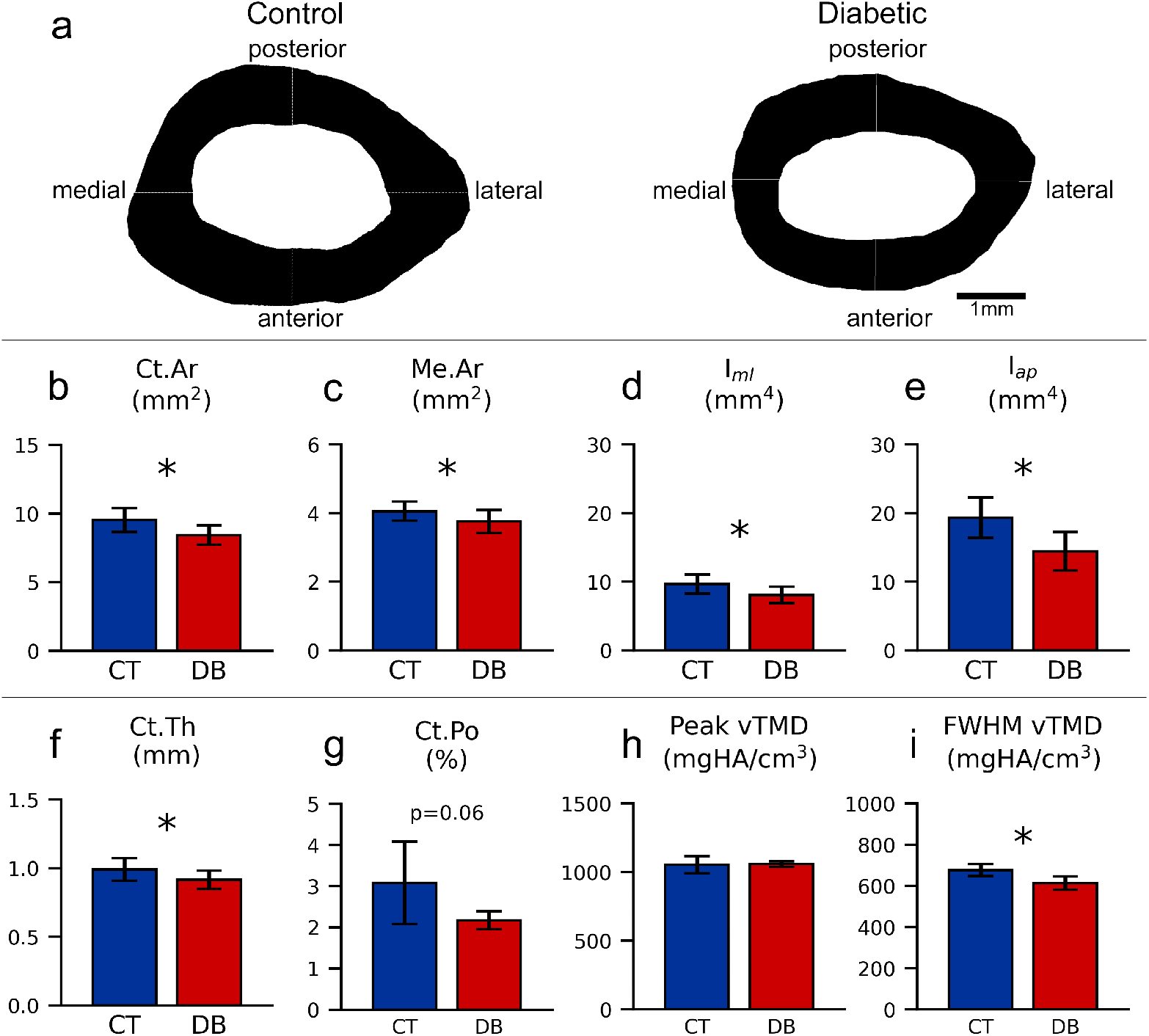
Geometry and microarchitecture measurements. Bone geometry and porosity obtained from microscopy and SRμCT are altered in diabetic (DB) rat bone compared to control (CT) rat bone. Optical images of bone fracture surfaces in diabetic (n=15) and control (n=12) rat femurs were (a) segmented to quantify geometrical changes between groups. (b) The cortical bone area of the diabetic group is significantly lower than the control group (−11.6%, p=0.002), as well as (c) the medullary cavity area (−7.3%, p=0.025). (d) The moment of inertia relative to the medial-lateral axis (−16.6%, p=0.005) and (e) the moment of inertia relative to the anterior-posterior axis (−25.3%, p=0.0003) were significantly lower in the diabetic group. (f) Cortical thickness was reduced with diabetes (−7.5%, p=0.02). SRμCT revealed that (g) cortical porosity was decreased by 30% (p=0.06) in diabetic rats (n=6) when compared to the control (n=4), whereas (h) the peak of volumetric tissue mineral density (vTMD) value (i.e., most common mineral level) remained unchanged between groups, with a 0.1% change in value (p=0.98). (i) The full-width at half-max (FWHM) of the vTMD distribution decreased significantly by 10% with diabetes (p=0.02). Data are given as mean ± s.d. Groups were compared with an independent t-test.

### T2DM compromises canal and osteocyte lacunar microstructure

To further evaluate the 30% reduction in cortical bone porosity with diabetes, we imaged and quantified canals and osteocyte lacunae at the micrometer scale using SRμCT. Canal morphology revealed prominent changes in the number of canals from control (n=4) to diabetic bone (n=6). Representative 3D images of control (Fig. 4a) and diabetic (Fig. 4b) bones show differences in the number of canals between the two groups, which were quantified with canal density (Ca.Dn) (Fig. 4c), the number of canals per bone volume. We measured a significant decrease of 31% in Ca.Dn when comparing control and diabetic rat bone (p=0.01). Although we measured marked changes in Ca.Dn, changes in the diameter of canals (Ca.Dm) (Fig. 4d) were not as prominent. We found a slight increase in Ca.Dm (3.4%) when comparing the control rat bones to the diabetic rat bones. Although this tendency for increased Ca.Dm can also be observed visually in Fig. 4b, the changes in diameter were not statistically significant (p=0.18).

**Fig. 4.**
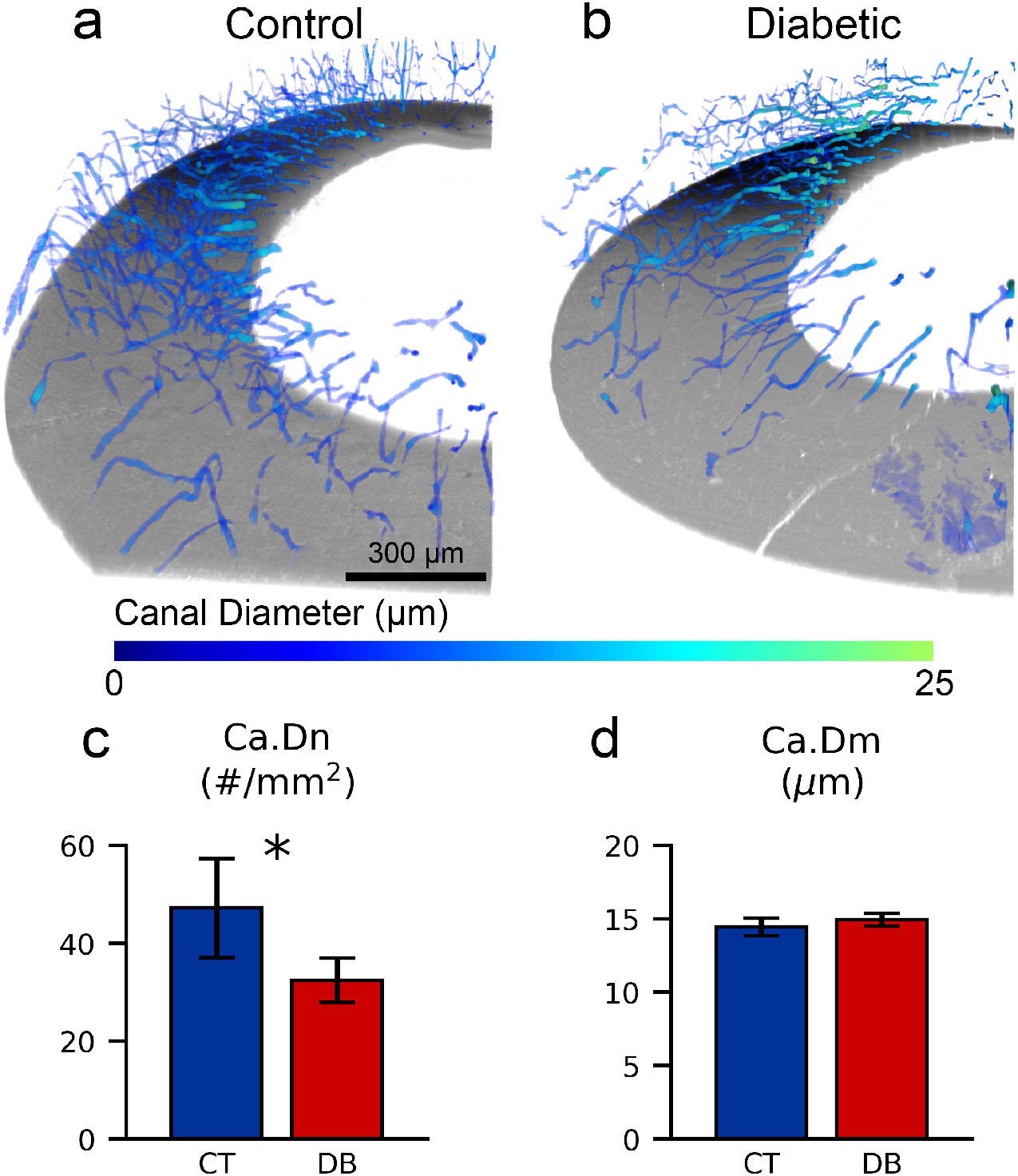
Canal distribution and size. Canal density and diameter in control (CT, n=4) and diabetic (DB, n=6) rats were measured using SRμCT. Decreased canal density and increased canal diameter are shown visually in representative SRμCT images for (a) control rats and (b) diabetic rats. (c) Canal density was significantly reduced by 31% in diabetic rats compared to control rats (p=0.01). Our analysis of (d) canal diameter revealed a slight increase in canal diameter of 3.4% in diabetic rats compared to control rats, though this change was not statistically significant (p=0.18). Data are given as mean ± s.d. Groups were compared with an independent t-test.

Similarly to the canal metrics, we also measured changes in the volume and density of osteocyte lacunae when comparing control (n=4) (Fig. 5a) and diabetic (n=6) (Fig. 5b) rat bones. Osteocyte lacunar density (Lc.Dn) (Fig. 5c), or the number of lacunae per bone volume, is greatly decreased in diabetic rats compared to the control (p=0.054). This 16% decrease in Lc.Dn is shown visually in Fig. 5a and b and was accompanied by a decrease in mean lacunar volume (Lc.V) (Fig. 5d). Lc.V was significantly reduced by 14% in diabetic bone (p=0.02). Through SRμCT, we found notable reductions in lacunar density and size with diabetes; we also measured concurrent decreases in the density of both canals and lacunae in ZDSD rats.

**Fig. 5.**
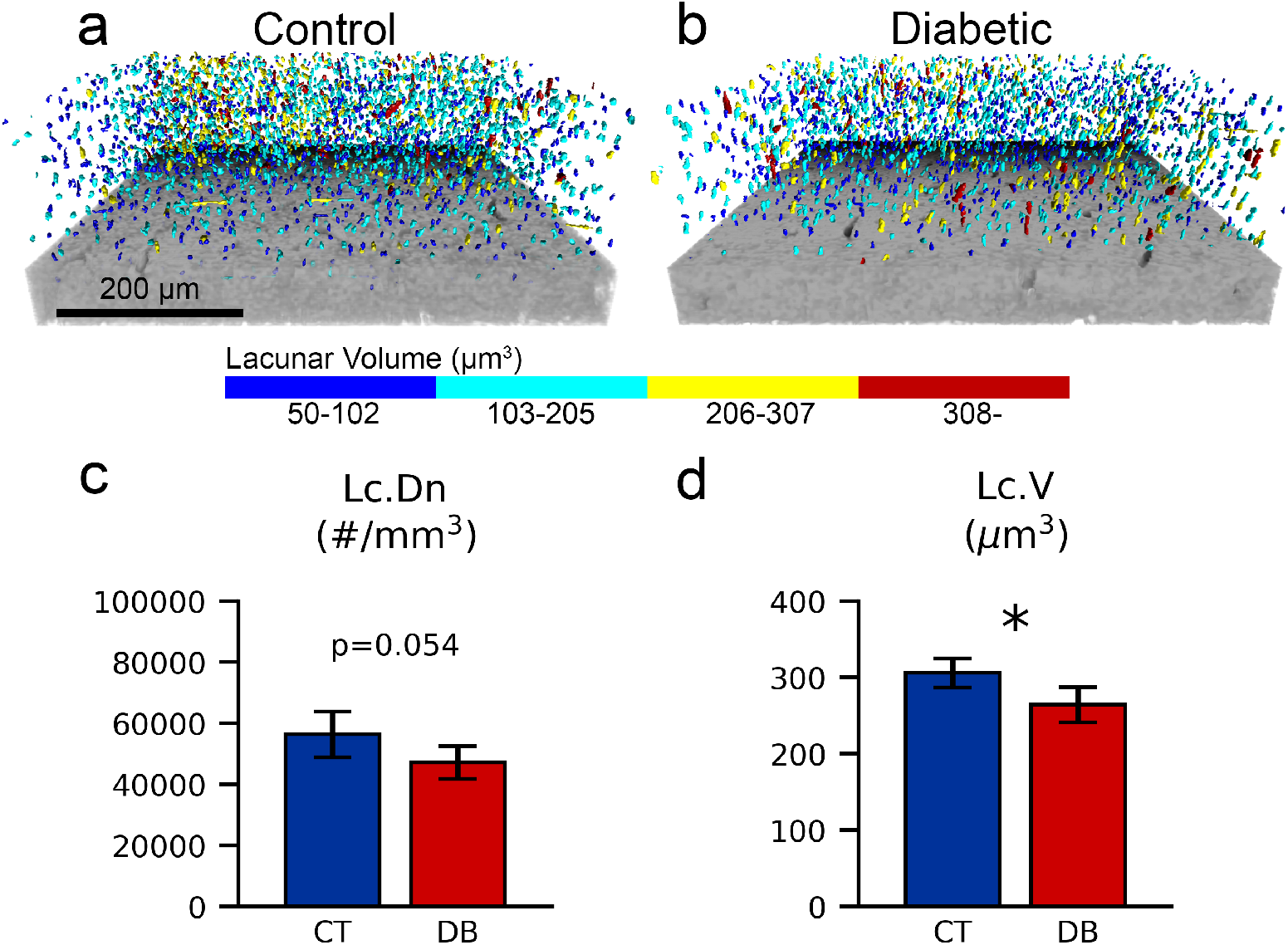
Lacunar density and volume. Lacunar density and volume were analyzed using SRμCT for control (CT, n=4) and diabetic (DB, n=6) rats. Representative 3D images of (a) control and (b) diabetic rat bone show visual changes in the number and volume of osteocyte lacunae. These visual differences are exemplified when quantifying (c) lacunar density, where a 16% decrease is observed in diabetic rats compared to the control rats (p=0.054). Additionally, (d) mean lacunar volume was concurrently decreased by approximately 14% in diabetic rats when compared to their controls (p=0.02). Data are given as mean ± s.d. Groups were compared with an independent t-test.

## Discussion

AGEs accumulation in T2DM is recognized as a significant contributor to diabetic bone fragility and loss of bone quality [13, 14, 53]. Several studies have shown that increased levels of bone AGEs restrict the collagen fibril deformation in T2DM, causing deficits in post-yield bone properties and energy dissipation [10, 15, 54]. However, the impact of T2DM on microstructural changes associated with bone remodeling and decreased toughness has not been studied yet. In this study, we investigated changes in microstructural features and porosity, measured by SRμCT, associated with loss of bone toughness and material properties, measured by *in situ* SEM fracture toughness and strength tests. Using a polygenic T2DM rat model, we found that nineteen-week-old diabetic rat femurs have half of the crack-growth toughness of control femurs. Crack-growth resistance to fracture in bone is conferred primarily by crack deflection around microstructural features. This loss of fracture resistance coincides with 1) a 15-30% reduction in canal and osteocyte lacunar density, and 2) a 13-15% reduction in post-yield properties (mostly post-yield strain). The decrease in osteocyte lacunar density, the void housing the main cells regulating bone remodeling, is also a sign of impairment in osteocyte cell viability and function, known to impact bone quality.

In this study, the diabetic rat phenotype closely mimics T2DM in humans; ZDSD rats exhibit low body weight, elevated fasting glucose, elevated levels of AGEs, and decreased bone resistance to fracture with the development of diabetes. The male ZDSD rat model was developed as a pre-clinical model of human diabetes closely resembling the human condition by having normal leptin signaling [55] and development of diabetes after skeletal maturity under normal diet [10, 43, 44, 56]. We found an 8% lower body weight in diabetic animals and a corresponding 7-12% significant decrease in whole-bone cortical geometry (e.g., cross-sectional area, cortical thickness, medullary cavity area). This is consistent with previous studies on ZDSD rats [10, 44] and follows weight loss seen in patients developing diabetes. Weight loss is a compensatory mechanism that some diabetic patients exhibit when developing the disease; the body fat is burned for energy to compensate for the inability of glucose to move from the blood to the cells. ZDSD rats also displayed a 44% increase in fasting glucose level (reaching hyperglycemic levels) associated with a 54% gain in AGEs content. Our glucose values match values found in the literature [10, 44, 57, 58] and resemble increases in blood glucose concentrations that occur in humans [55, 59, 60]. As expected, hyperglycemia was associated with a fast and significant increase in AGEs content in ZDSD rats, similar to the pentosidine increase measured by Creecy et al. in 22-week-old ZDSD rats [10]. AGEs accumulation in T2DM has been shown to be a key contributor to loss of bone fracture resistance and bone quality in diabetic bone [10, 15, 54].

The novel finding of this work is that diabetes significantly decreases canal and osteocyte lacunar density and total volume. We measured a 31% and 16% decrease in canal and lacunar density, respectively, with a 14% reduction in lacunar volume. Combined, this resulted in a 30% decrease in cortical porosity. This significant decrease in canal density observed in ZDSD rats is consistent with other diabetic and hyperglycemic models recorded in the literature [61, 62]. Such a decrease in canal density will impair extrinsic toughening mechanisms that use microstructural features to dissipate energy mostly via crack deflection [41, 63]. More specifically, the decrease in canal density indicates a decrease in osteon density, which will reduce the potential of crack deflection on cement lines around osteons [42, 64]. Moreover, decreased osteocyte density is known to result in an accumulation of microdamage and microcracking [21, 22], which will also alter extrinsic toughening mechanisms. Our *in situ* SEM fracture toughness results showed that T2DM induces a 55% reduction (p=0.09) in crack-growth fracture resistance, which can be associated with these microstructural changes. In addition, we found a significant reduction in post-yield properties, mostly through post-yield strain (13% decrease in strain to failure), which is a well-known consequence of AGEs increase with diabetes. Indeed, an increase in AGEs content restricts collagen’s ability to deform, which ultimately limits bone’s post-yield properties and work to fracture [15]. This loss of material properties and plastic deformation limits intrinsic toughness, which makes a crack easier to initiate and propagate [36]. Even though this is not the main mechanism contributing to crack-growth fracture resistance, it might have participated in the loss of fracture resistance in addition to microstructural alteration with diabetes. The decrease in crack-growth toughness exhibited by ZDSD femurs is paralleled by other studies [44, 65–67]. However, Creecy et al. reported a significantly decreased toughness only with the advancement of the disease from 16 to 29 weeks, not against the control, using a similar single-edge notched toughness test as the one performed in this current work [10]. Because extrinsic toughening mechanisms are thought to be more effective than intrinsic mechanisms [35], we attribute the decrease in crack-growth toughness seen in ZDSD rats to primarily decreased canal density, but also to loss of material properties associated with AGEs accumulation. Indeed, both AGEs accumulation and microstructural changes together build the whole picture of impaired toughness in T2DM bone.

In addition to their role in toughening mechanisms, canals are also vital for nutrient transport to the bone cells [68]. The concurrent change in canal and lacunar density may suggest that vascularization is decreased, indicating a reduction of bone cell activity. Osteocyte lacunae give insight into the number and the state of osteocyte cells. Because these cells play an important role in the regulation of skeletal homeostasis, a greater mean lacunar volume may imply increased remodeling [69], whereas a small volume can represent a trend toward osteocyte apoptosis. T2DM has been shown to affect osteocytes, resulting in their dysfunction and changes in lacunar density and size. The nature of these changes is inconsistent in the literature. Villarino and colleagues report significantly decreased lacunar density in streptozotocin (STZ)-induced diabetic rats [70], while others report modest, non-significant decreases [61]. Some report an increase in lacunar density [71]. These variable changes in lacunar density indicate that the mechanisms influencing osteocyte lacunae number may not be solely due to hyperglycemia, which has been consistently elevated in all the noted studies. Although the ZDSD rat phenotype used in our current work shows decreased lacunar density, the effects of STZ injection and high-fat diet (HFD) on lacunar density may be different among the myriad animal models. Changes in canals and osteocyte lacunae might also be related to bone marrow adipose tissue and adipocytes [9, 72–75] or increased inflammation [76]. Overall, these varied results indicate the need to investigate mechanisms affecting the lacunar density and bone remodeling in T2DM, especially in regard to the responses of differing models.

Finally, an explanation of the lacunar and canal density decrease could come from the accumulation of AGEs in T2DM. Several studies showed that AGEs affect bone cells, such as osteoblasts, osteocytes, and osteoclasts [20, 77, 78]. One repercussion of AGEs accumulation is the increase in sclerostin, an osteoblast inhibitor [79], and the decrease in receptor activator of nuclear factor-*κ*B ligand (RANKL) [25], a key factor in osteoclastic differentiation and action [80]. These changes in sclerostin and RANKL show that AGEs alter bone remodeling. Because AGEs accumulation is due to an imbalance between AGEs formation and resorption, once created, AGEs exacerbate their own accumulation by inhibiting their resorption. The reduction in bone remodeling is expected from the change in canal and osteocyte lacunar density. The reduction in osteocyte lacunar density may show that multiple osteocytes die and that the voids around these cells were filled by the mineral matrix. This mechanism is called micropetrosis [81]. Moreover, since the mean lacunar volume is decreased, we can interpret that the osteocyte activity is decreased because remodeling starts from the removal of the perilacunar matrix in healthy bone [69]. We did not directly measure the osteocyte activity, but thanks to the quantification of osteocyte lacunae, we managed to show that T2DM reduces bone remodeling and interpreted that AGEs are responsible for this reduction. Targeting AGEs accumulation or AGEs effects on bone cells could limit the decrease in toughness taking its origin at the microscale.

Although the present study succeeds in assessing the mechanical and microstructural properties of diabetic rats, it also possesses some key limitations. Firstly, although our rats were diabetic for three weeks, they did not develop severe diabetic complications, thus reducing the generality of our conclusions. We would advise measuring the HbA1c, characteristic of the level of blood sugar level over a period of three months, for better insight of the disease advancement. This measurement would be helpful, especially since we showed that AGEs content or hyperglycemia might not be enough to estimate the effect of diabetes on biological and microstructural features, as seen in STZ and HFD rats, for example [61, 70]. However, one repercussion of diabetes is AGEs accumulation, and our rats indeed displayed this accumulation. Older rats would likely exacerbate this accumulation and intensify our findings. Second, there is no ASTM standard to calculate stress intensity factors from the toughness test for a rat bone cross-section. We followed the method published by Ritchie et al. [47] on small animal bone testing. This method assumes a circular cross-section, but since rat femurs are more elliptical, it induces approximately 17% uncertainty on the stress intensity factor. Moreover, the solution for *J_pl_* is valid for 20 > Rm/t >5, when in rat femur is generally around 2 (1.74 in our case), explaining the p-value of the crack-growth toughness. Finally, we aimed to have eight samples per group to have higher statistical power, but due to equipment constraints, most of our tests have n=4-6. This induces more variability in our findings, but our statistical power is still ranging between 60% for lacunar density to 96% for canal density.

To our knowledge, we have revealed for the first time that T2DM detrimentally impacts microstructural features, such as lacunae and canals, responsible for loss in toughness and material properties of diabetic bones. Two mechanisms can explain the decreased fracture toughness in ZDSD rats: (1) AGEs accumulation impairing bone material properties via loss of deformation, and (2) alteration of bone remodeling and cell function affecting bone microstructure. Since AGEs seem to be one of the key proteins altering bone integrity, reducing factors that induce AGEs and targeting AGEs resorption would improve bone quality. Microstructural feature analysis on diabetic rat models with treatment that inhibits AGEs would give insight into the role of AGEs and could help reduce the fracture risk in the diabetic population.

## Acknowledgments

This work received support from the National Science Foundation under NSF CAREER Grant CMMI 2045363. This work also used resources from the Advanced Light Source at beamline 8.3.2., a U.S. DOE Office of Science User Facility under contract no. DE-AC02-05CH11231.

## Author contributions

C.A. designed research; W.W., Y.O., and K.M. performed research; W.W. and Y.O. analyzed data; W.W., Y.O., and C.A. wrote the manuscript. All authors have given approval for the final version of the manuscript.

## Competing interests

The authors declare no competing interests.

## Notes

### Competing Interest Statement

The authors have declared no competing interest.

